# Time- and frequency-resolved decomposition of beta brain activity reveals two functionally distinct beta bands

**DOI:** 10.1101/2025.11.10.682052

**Authors:** Sara Korkealaakso, Christine Ahrends, Linda Niemelä, Diego Vidaurre, Hanna Renvall, K. Amande M. Pauls

## Abstract

Oscillatory brain activity in the beta (13-30 Hz) range plays a central role in sensorimotor and other healthy brain processes *and is a potential disease biomarker*. However, even in the well-studied sensorimotor system, inconsistencies remain concerning beta’s differential role in maintaining and terminating movement. Emerging evidence suggests that beta comprises two frequency ranges, a lower (<20 Hz) and a higher (>20 Hz) subrange. Moreover, beta occurs in transient ‘events’ which correlate with perception and action. We here use human whole-brain magnetoencephalography (MEG) recordings and a Hidden Markov Model based analysis approach to discover whole-brain beta-band activity during rest and a naturalistic motor task in a data-driven manner. We successfully delineate two anatomically and functionally distinct beta band components which coexist throughout the neocortex. Low-beta (<20 Hz) events are very rare, high amplitude events, whereas high-beta (>20 Hz) is common but lower in amplitude. Both bands show state-specific anatomical distribution and modulation during a motor task: Low-beta is entirely absent during movement, whereas high-beta is only partially suppressed and reoccurs during postural maintenance. During post-movement beta rebound, low-beta event probability increases threefold, accompanied by strong low-beta event synchronization. A third state, probably corresponding to gamma activity, is active during task execution. Beta rebound occurs via a directed, sequential shift from gamma/active state via high-beta state to low-beta state. Our results provide strong experimental evidence for the coexistence of two functionally and anatomically distinct beta bands, and provide insights into their role in movement initiation, maintenance, and termination.

**Short abstract:** The brain’s beta band activity is modulated by sensorimotor processing, but inconsistencies remain concerning beta’s role in maintaining and terminating movement. Mounting evidence points towards two beta bands with distinct functional roles. We use human whole-brain magnetoencephalography and a data-driven Hidden Markov Model based analysis approach to detect whole-brain beta-band activity. We delineate two anatomically and functionally distinct beta band states: a rarely occurring low-beta (<20 Hz) and a common high-beta (>20 Hz) state. During movement, low-beta is completely absent, whereas high-beta is only partially suppressed. During post-movement beta rebound, only low-beta increases significantly, accompanied by pronounced low-beta synchronization. Activity in gamma state increases during movement. Beta rebound occurs via a sequential shift from gamma via high-beta to low-beta state. Our results provide strong experimental evidence for the coexistence of two functionally and anatomically distinct beta bands and provide insights into their role in movement initiation, maintenance and termination.

## Introduction

Oscillatory activity in the so-called beta range (13-30 Hz) is one of the most salient rhythms that the human brain generates, and it is found also in other species, e.g., primates ^1–3^ and rodents ^2,4^. Beta rhythm was first described in the sensorimotor cortex ^5^, but there is mounting evidence for a functional role of beta activity beyond strictly sensorimotor processing ^6^, including working memory ^7,8^ and executive control ^9^. In addition to the sensorimotor cortices, beta-range activity can be observed, e.g., in the frontal cortex ^7,9^, as well as in the basal ganglia ^4,10,11^. Furthermore, beta oscillations have been demonstrated to synchronize over cortical and subcortical networks ^12–14^, suggesting a role in long-range connectivity.

During sensorimotor processing ^15^, cortical beta activity is modulated in a movement-dependent manner: It is suppressed during brief movements ^16,17^, but increases during postural maintenance ^18–20^ and stabilizes with stable posture ^19,20^. Cortico-muscular coherence is also predominantly observed in the beta band ^18,20^. After movement completion, beta activity increases ^16^: This so-called post-movement beta rebound is modulated, e.g., by kinematic errors ^21^ and movement cancellation ^9,22,23^. The rebound has been interpreted as an indicator of movement outcome processing ^24^, or an index of confidence in predicting motor outcome ^25^. Thus, beta modulation depends considerably on the type of movement studied.

There is emerging evidence that the beta band comprises two separable frequency ranges, a lower (< 20 Hz, low-beta) and a higher (>20 Hz, high-beta) subrange ^26,27^. It has been proposed that low-beta has an “anti-kinetic” role ^28,29^, while high-beta might reflect attention or sensory cue anticipation ^3,29–31^. This dissociation into two distinct beta bands may explain some of the inconsistencies in experimental findings concerning task-related beta behavior, including the discrepancy between brief and sustained movements. Moreover, beta activity is not continuous, but occurs in transient ‘events’ whose probability varies with task phase ^1,2^, peaking after movement completion to produce the beta rebound phenomenon. Occurrence of transient beta events predicts the detection probability of tactile stimuli ^32,33^. Human cortical sensorimotor beta events at rest are highly stable ^34^ and show heritable features ^35^. Beta activity and beta events are also of great clinical relevance, as they are altered, e.g., in Parkinson’s disease ^36,37^ and stroke ^38,39^, where beta correlates with hand function ^38^.

Thus, the beta rhythm and its sub-bands - with their spontaneous as well as task-induced fluctuations over time - are thought to play a crucial role in many healthy brain processes and are a promising potential disease biomarker. Nevertheless, even for well-studied healthy motor processing, the beta rhythm’s differential modulation, with suppression during short, phasic movement and relative increase during force maintenance, is not fully understood. Analyses accounting for both the time-resolved nature of the beta rhythm, as well as its potential segregation into two separate frequency bands, can enrich the information obtainable from electrophysiological data and help to resolve inconsistencies.

To address this, we used an analysis approach based on the Hidden Markov Model (HMM). Essentially, the HMM framework segments timeseries data into a sequence of a finite number of states. Vidaurre et al. have previously implemented a HMM-based approach for segmenting multivariate electrophysiological signal time series into states characterized by their unique spectral properties ^40,41^. We here adapt this method to decompose spectral oscillatory behaviour at the level of individual parcels and apply it to human MEG data during rest and a naturalistic motor task, simultaneously estimating from the data both when specific events occur (temporal properties) and what is their dominant frequency (spectral properties).

We can decompose beta activity into two distinct beta states, a rarely occurring lower (13-20 Hz) and a more frequently occurring higher (20-30 Hz) beta frequency state. We show that the two beta states are differentially modulated by task, with total suppression of low-beta throughout motor execution versus partial suppression of high-beta during motor task initiation and termination, but partial recovery during force maintenance. Low-beta accounts for the very pronounced, highly synchronized beta rebound response. During the post-movement rebound, there is a sequential shift from gamma/active state via high-beta to low-beta state. Our findings strongly support the existence of two functionally distinct beta bands and provide detailed insights into their role and interplay in movement initiation, maintenance, and termination.

## Results

Resting state MEG data was recorded from 42 healthy adults (mean age 45.7 +/- 14.8 years, range 18-78 years, 15 females). A subset of the subjects (n=22, mean age 48.5 +/- 15.2 years, range 18-78 years, 10 females) also performed a cued motor task, squeezing a ball with their left hand for two seconds (**figure 2D**). The analysis pipeline is depicted in the project overview in **figure 1**; further details can be found in the Methods section. Briefly, band-pass (13-30 Hz) filtered MEG data was first transformed to source space using the minimum norm estimate algorithm implemented in MNE-python ^42,43^. Source time series dimensionality was reduced to n=448 parcel time series (using the aparc_sub parcellation ^44^) via principal component analysis of all voxel time series within one parcel, resulting in one principal component time series ^42^ per parcel.

**Figure 1:**
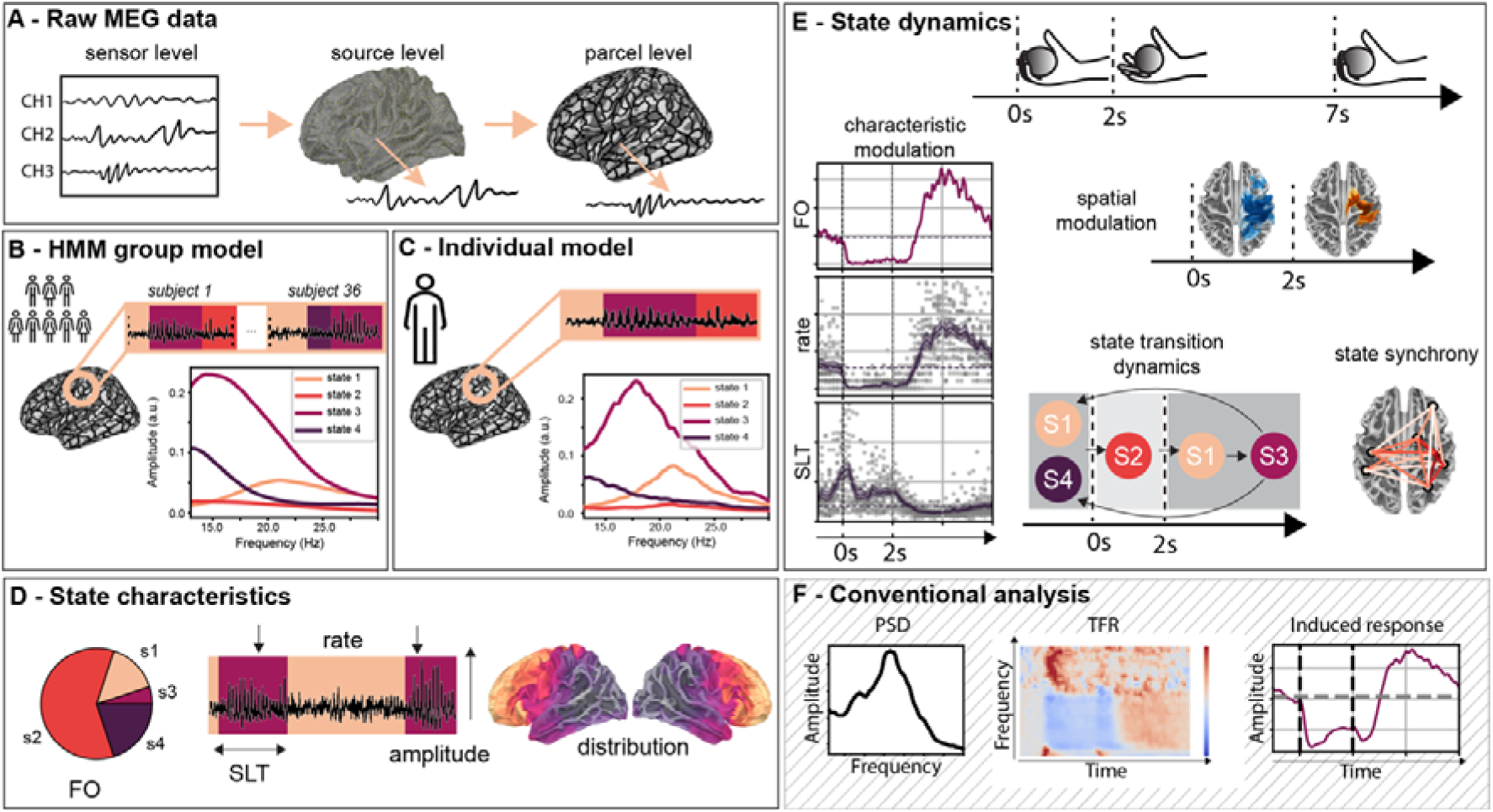
Study overview and illustration of HMM beta event analysis pipeline. **A**. Raw MEG time series data is band-pass filtered (13-30 Hz) and transformed from sensor to source to parcel level using MNE python ^45^. **B.** Using the HMM-MAR toolbox time-delay embedded implementation ^46^, a group model is established on a reference group, **C.** and then fitted to individual subjects to obtain individual subjects’ state time series and spectral characteristics per brain parcel. **D.** Binary state time courses (Viterbi paths) and probability time courses from these individual HMMs, together with the MEG time series, are used to calculate event characteristics: State lifetime (*SLT*) describes the length of each state visit. Fractional occupancy (*FO*) describes the share of time spent in a given state. Event amplitude (*EA*) describes the amplitude of the signal amplitude envelope during each state visit. Events per second (*rate*) describes how many events occur per second. In addition, anatomical distribution of these parameters can be inferred. **E.** Dynamic modulation of state characteristics in relationship to the task and its anatomical distribution can be examined for different states, as well as synchronization across areas within state, and dynamics between states and their timings, enriching the information that can be obtained from the MEG data. **F**. Conventional MEG data analysis (power spectral density, time-frequency analysis, band specific induced responses). HMM: hidden Markov model, PSD: power spectral density, TFR: time frequency representation

**Figure 2:**
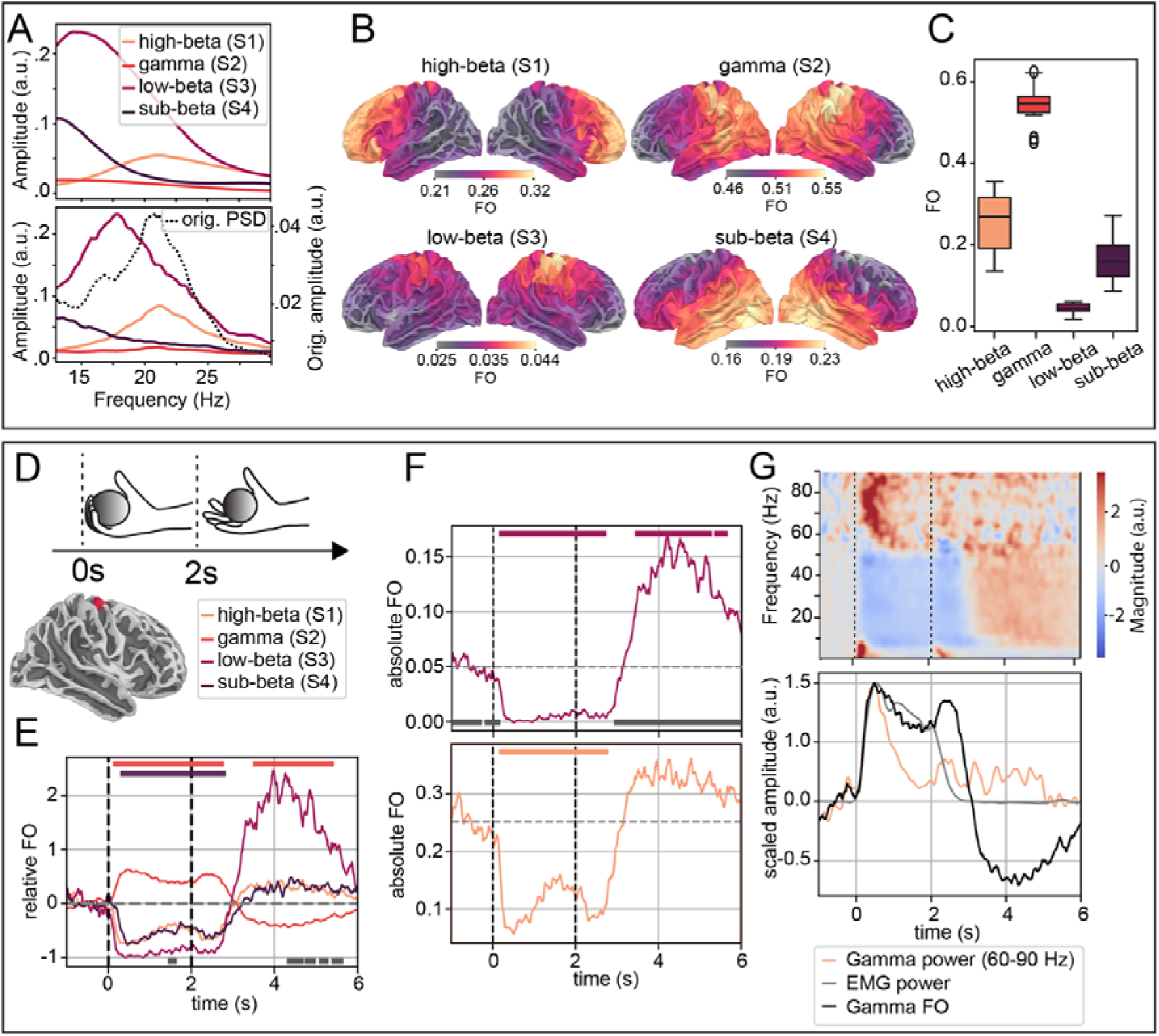
Beta range spectra for the four states obtained with the HMM method **A.** (upper panel) from the group HMM model and **A.** (lower panel) in a representative parcel of an individual subject (contralateral M1). The black dotted curve in **A.** (lower panel) shows the power spectral density curve obtained from conventional (multitaper) spectral analysis (same individual and parcel). **B.** Fractional occupancy (FO) distribution across all cortical parcels during resting MEG brain activity (eyes open) for the four observed states, showing distinct anatomical distribution and maxima. **C.** Boxplot illustration of resting state FO distribution, per state and across all parcels, showing very distinct FO patterns, with very rare low-beta and more common high-beta and ‘gamma’ states. **D.** Schematic of the motor (ball squeeze) task. **E.** Task-evoked FO relative to the baseline interval (all four states), and **F.** absolute task-evoked FO change (for the high and low-beta states) during the motor task (contralateral hand area M1 parcel, localization shown in brain inset in E). Significant difference from baseline is indicated by same-colored bar (top); in E, the grey bar (bottom) indicates significant difference between low and high-beta state; in F, the grey bar (bottom) indicates significant difference from 0 (non-parametric cluster-level paired t-test for temporal data, cluster threshold p<0.001, significance threshold p<0.001, for further details see Methods section). Horizontal dashed lines indicate the baseline level at rest, while vertical dashed lines at 0 and 2 seconds correspond to the ‘go’ and ‘stop’ cues. **G.** (*upper panel*) Conventional time-frequency representation of movement-related oscillatory MEG power; (*lower panel*) comparison of scaled (baseline interval set to zero, curve scaled to maximum of 1) 60-90 Hz MEG signal amplitude (orange), EMG power (grey), and gamma state FO (black) show a simultaneous increase. Gamma FO is maintained more steadily than gamma band power, following relative EMG power. At movement termination, gamma FO shows a transient increase followed by a relative suppression coinciding with the return of EMG power to baseline.

The HMM was first fitted at group-level using resting data from 36 subjects and concatenating all subjects and parcel timeseries, resulting in a common group model. This group model was fitted again to each individual subject and parcel, for both rest (all subjects) and motor task (n=22), giving an individual state time series and spectral estimate for each subject and parcel, per task ^41^. This so-called dual estimation (i.e., first estimating parameters at the group-level, and then estimating individuals’ parcel-level versions of group parameters) allows the spectral description to vary slightly between individuals and brain areas, while maintaining a common group reference, ensuring comparability. From the obtained state time series, we characterised the proportion of the time course taken up by each state (fractional occupancy, FO) and its anatomical distribution (**figure 1D**) at rest, and how these (FO and its anatomical distribution) are modulated by task, as well as state synchronization behaviour and state dynamics (**figure 1F**). We also derived the duration of state visits (state lifetimes, SLT), event amplitudes (EA), and the number of events per second (rate), using the state time series and/or the raw MEG data (**figure 1D**). This was conducted in a data-driven approach, i.e., we did not impose separate bands on the data but rather aimed to observe which frequencies naturally emerge from the data. Given our hypothesis about distinct low- and high-beta activity, we selected the HMM solution with the smallest number of states that captures this distinction —here four.

### Two distinct beta range states

First, we investigated the states’ spectral properties. Two distinct beta states, a low (peak < 20 Hz) and a high-beta state (> 20 Hz) (**figure 2A, state 1 (high-beta) and 3 (low-beta)**) were observed in all subjects. In addition, we found one state with aperiodic, 1/f -like spectral properties in the beta range. This likely represents the residue of oscillatory activity below beta range remaining after the band pass filtering applied here (here called ‘sub-beta state’ based on its anatomical distribution, see **figures 2A and 2B, state 4**), and one state with very little periodic activity in the beta range (**figure 2A, state 2**). We will refer to this state as ‘active’ or ‘gamma’ state, due to its behavior during the motor task (**figure 2E**) as well as its modulation with respect to high-gamma power and EMG power during the motor task (**figure 2G**).

### States have a distinct, state-specific topography but co-exist across the entire cortex

Next, we investigated states’ anatomical distribution across the cortex. Both beta states co-exist throughout the entire cortex, but their likelihood of occurrence (fractional occupancy, FO) varies with cortical area. The low-beta state is most likely to occur in parietal and posterior frontal (sensorimotor) areas surrounding the central sulcus, whereas the high-beta state has the highest probability of occurrence in the frontal cortex (**figure 2B).** The gamma/active state has a posterior frontal and parietal topography with a maximum around sensorimotor areas at rest, while the sub-beta state has its maxima in the temporal and occipital areas. States’ FO distributions are overlapping (**figure 2B**), indicating that all four states co-exist across the whole cortex.

### High-beta is common, while low-beta is very rare

While the high-beta state is common (resting FO 21-32%), the low-beta state has a very low occurrence rate (resting FO 2-4%, **figure 2C**). Of the four states, the gamma/active state is the most frequently observed.

### Low-beta is suppressed entirely during movement and underlies the post-movement rebound

Using individual state time series, we next analyzed how movement affects oscillatory behavior. States’ fractional occupancy is modulated by task (**figure 2E**): During sustained movement, the low-beta state is suppressed entirely in contralateral M1 (**figure 2F, upper panel**). In contrast, the high-beta state shows a significant, but not complete suppression associated with the initiation and termination cues for the movement, interspersed with a period of partial recovery during force maintenance (**2F, lower panel**). The beta suppression is followed by a post movement rebound after cessation of movement, which is significant only for the low-beta state, with a threefold FO increase compared to baseline (**figure 2F, upper panel**). The gamma/active state is modulated in the opposite direction to the high-beta state’s behavior: it increases when high-beta decreases and is suppressed during the beta rebound (**figure 2G)**.

### Modulation occurs in brain areas associated with movement control

State occupancy modulation localizes to anatomical areas involved in movement-related processing: Significant task-related modulation of FO occurs in a classical network of both contralateral and ipsilateral areas relevant for motor processing **(figure 3C)**, in a state- and task-phase specific manner and with good signal-to-noise ratio. Task-related modulation of FO differs both in pattern and magnitude between ipsilateral and contralateral cortical areas (**figure 3A**). Areas with significant FO modulation include contralateral/ipsilateral M1 and S1, contralateral SMA/PMd, contralateral prefrontal cortex (PFC) and contralateral posterior parietal cortex (PPC). All contralateral areas show a typical sustained beta suppression and rebound pattern. The suppression is biphasic for both beta bands in ipsilateral areas (iM1, iS1, **figure 3A and 3C**), as well as in cPFC, cSMA and cPPC for the high-beta band. Modulation coincides temporally with the initiation and release of ball squeeze (change in motor state), but is not maintained during force maintenance. Importantly, ipsilateral M1 and S1, as well as contralateral PFC show significant modulation non-contiguous with contralateral M1 and S1, speaking against source leakage from sensorimotor areas. The active/gamma state shows increased FO in contralateral motor areas during movement, and a relative suppression during the rebound (**figure 3C**). Thus, we can decompose activity in different frequency bands, showing distinct modulation for low- vs. high-beta band in different anatomical areas, with opposite behavior for beta and active/gamma frequency bands.

**Figure 3:**
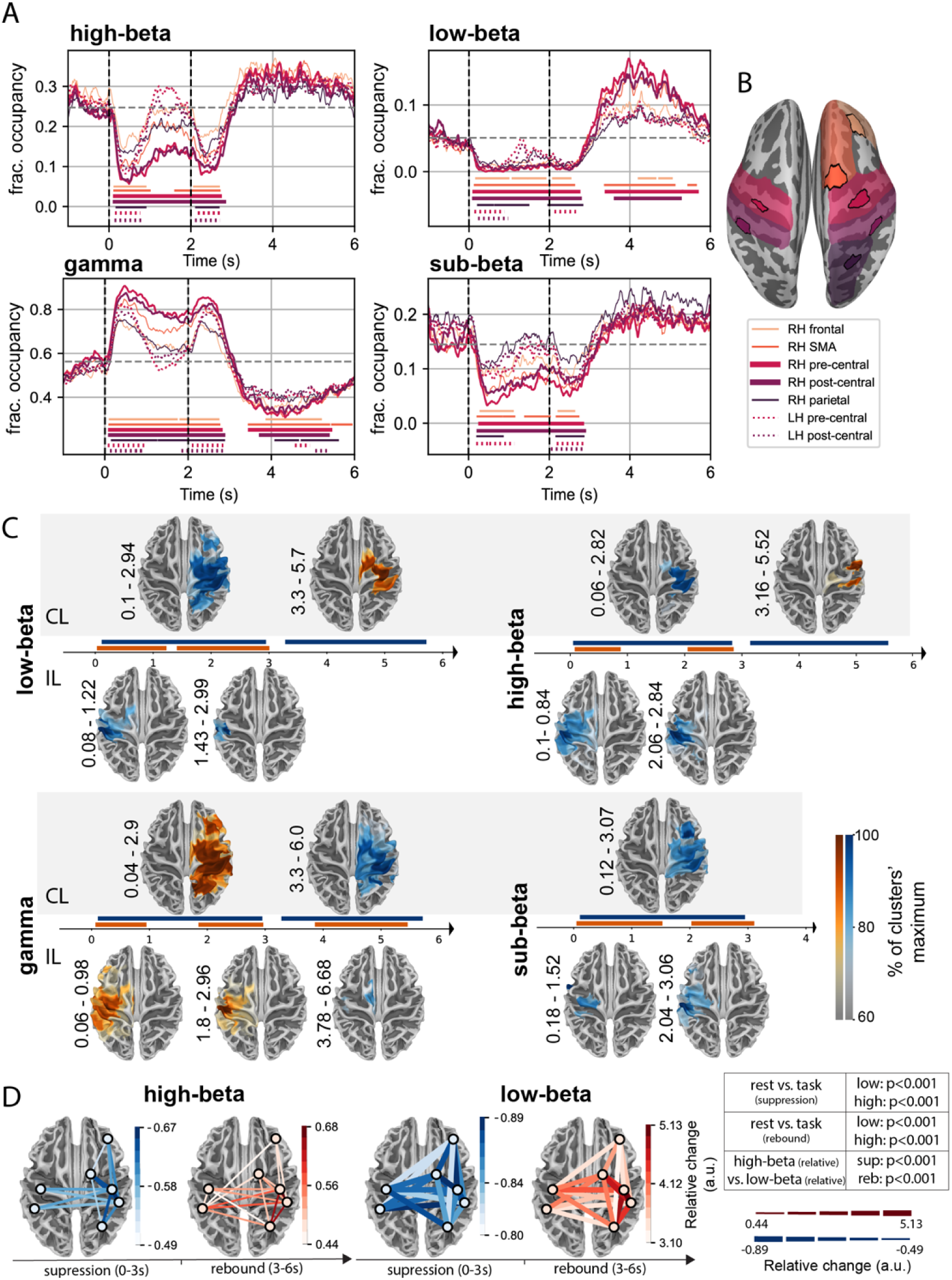
Task-induced modulation of state FO and state synchrony. **A.** Task-induced FO modulation profiles for all four states in the ipsi-(dashed lines) and contralateral (solid lines) cortical areas with significant task-related FO modulation. Significant differences from baseline are indicated by the colored bars at the bottom, matched to the curves in color and pattern (non-parametric cluster-level paired t-test for temporal data, cluster threshold p < 0.001, significance threshold p< 0.001). **B.** Anatomical rendering of analyzed parcels (labels given in the legend), (parcels color-matched to the lines used in A). **C**. Clusters with statistically significant FO modulation during the motor task compared to the baseline interval (-1-0 s), shown for all states (non-parametric cluster-level paired t-test for spatio-temporal data, cluster forming threshold p < 0.001, significance threshold p < 0.001). The horizontal axis represents time (s); cluster significance times (s) are shown on the vertical axis for each cluster. CL - contralateral, IL-ipsilateral; blue = relative decrease (i.e., suppression), red = relative increase (i.e., rebound) in FO. The colors are scaled within each state to the state’s cluster maximum, thus representing variation within state. FO modulation occurs in a typical network associated with execution of a simple motor task. Both beta states show a suppression-rebound behavior, while ‘gamma/active’ state shows the reverse behavior, with task-related increases and suppression after task completion. **D**. Coincidence of events across different cortical areas for high-beta (left) and low-beta (right) states compared to the respective coincidence rate during rest, shown separately for the beta suppression and rebound phases. Since the strength of the synchrony modulation differed between low-beta and high-beta states by up to factor of 10, synchrony modulation is indicated by two different scales, once by line color (separate scales for high-beta and low-beta separately) and once using line thickness to visualize this difference between the (scaling identical for high-beta and low-beta), blue – decrease, red – increase. State synchrony modulation is much more pronounced for the low-beta state, and both event desynchronization and synchronization are significantly stronger for low- than for high-beta band. Both event desynchronization during beta suppression, as well as event synchronization during beta rebound are significant compared to rest for both beta bands (Mann-Whitney U-test, significance levels given in the table).

### Low-beta events are highly synchronized during the beta rebound

Since beta is believed to support long-range synchronization of brain activity, we investigated low- and high-beta states’ synchronization behavior using event coincidence, i.e., by assessing whether events occur simultaneously in pairs of areas, compared to resting state (see **Methods** for details). State synchrony in the sensorimotor network was significantly reduced in both beta states compared to rest during beta suppression and significantly increased during beta rebound (**Figure 3D**). Both desynchronization during the suppression, as well as synchronization during the rebound, are more pronounced for the low-beta state, with an almost 5-fold increase in beta event synchrony for the low-beta state during rebound, compared to a 1.7-fold increase for the high-beta state. Hence, the beta rebound is dominated by a pronounced increase in low-beta band activity that is synchronized across areas within the motor network.

### State transitions are modulated by task and occur in a sequential manner during post movement beta rebound

Using the discrete state time series obtained from the HMM based approach, we investigated transition dynamics between states in relationship to movement, which is not possible with conventional time series analysis. State transitions were grouped according to state pair and direction and visualized using a sliding window average. This analysis was carried out for contralateral M1. Movement initiation following the go cue is accompanied by significantly increased transition rates to gamma/active state from both high-beta and sub-beta states (**figure 4, 2^nd^ column**). After the stop cue, on the other hand, we observe significantly increased transition rates from gamma/active state to high-beta state (**figure 4, 1^st^ column**), and from high-beta to low-beta state, coinciding with the post movement beta rebound (**figure 4, 3^rd^ column**). The increase in transitions from active/gamma to high-beta state precedes the increase in transitions from high-beta to low-beta. Thus, after movement termination, cortical oscillatory activity changes in a directed, sequential manner from gamma/active through high-beta to low-beta oscillatory state. Furthermore, there were virtually no transitions between gamma/active and low-beta state and vice versa, and the observed transitions did not meet the expected transitions particularly during rebound (chi-square test, suppression: (from gamma: *p* = 0.008, from low-beta: *p* = 2.246e^-11^) rebound: (from gamma: *p* = 8.573*e*^-14^, from low-beta: *p*= 1.738*e*^-25^)).

**Figure 4:**
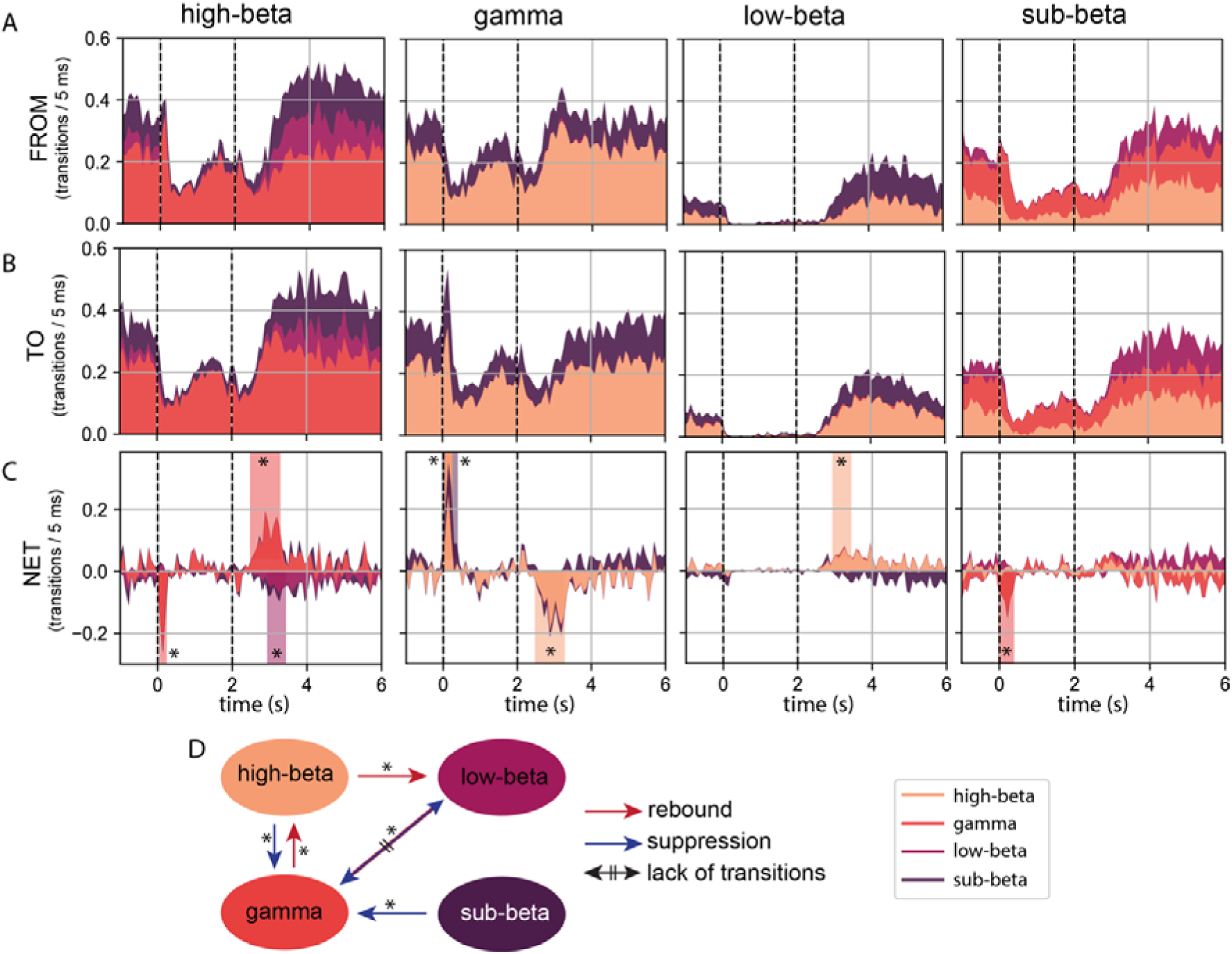
State transition dynamics during the motor task. Columns correspond to each state, and stack plots show transitions **A.** away from the state, **B.** transitions to the state, and **C.** net transitions, indicating the dominant direction of the transition flow (+ towards the state, - away from the state). Background shading indicates periods with significant net transitions to/from the state, the shading colour indicates from/to which state the transition occurs (MNE’s non-parametric cluster-level pared t-test for non-spatial data, compared to baseline interval (-1,0 s), cluster threshold p < 0.05, significance threshold p < 0.001). Dashed vertical lines indicate movement task go and stop cues (at 0 and 2 seconds). **D.** Schematic summary of the significant transition dynamics between the states and their direction during suppression (red arrows) and rebound (blue arrow).

### State characteristics are modulated by movement and can carry independent information about neural processing

So far, we have considered states’ FO and its modulation and synchronization with movement, but other state characteristics, such as event rate and amplitude, and their modulation may also carry significant information. During the motor task, event rate decreases significantly for all four states (**supp. figure 1B**) but increases significantly only for low-beta during the beta rebound (**supp. figure 1B**, third row). State lifetimes are significantly increased for low-beta during the post-movement rebound, and significantly decreased for the active/gamma state (**supp. figure 1A**). Event amplitude is reduced for all states during movement and increased during rebound (**supp. figure 1C**). While gamma/active state FO (**figure 2E**) and state lifetimes (**supp. figure 1A, second row**) are reduced during the post-movement period, event amplitudes are increased (**supp. figure 1C, second row**), suggesting that different state characteristics can carry distinct and orthogonal information about neural processing.

During rest, event rate **(supp. figure 2C)** and state lifetime **(supp. figure 2A)** show state-specific anatomical distribution patterns: While gamma state lifetimes are longest in areas surrounding the central sulcus, its rates are highest in frontal areas, again illustrating that different characteristics can carry orthogonal information. Amplitudes for all states are highest in the sensorimotor cortices (**supp. figure 2B).** Thus, amplitude information in signal analysis (as used in conventional power-based analysis) can bias results towards areas surrounding the central sulcus.

**Supplementary figure 1:**
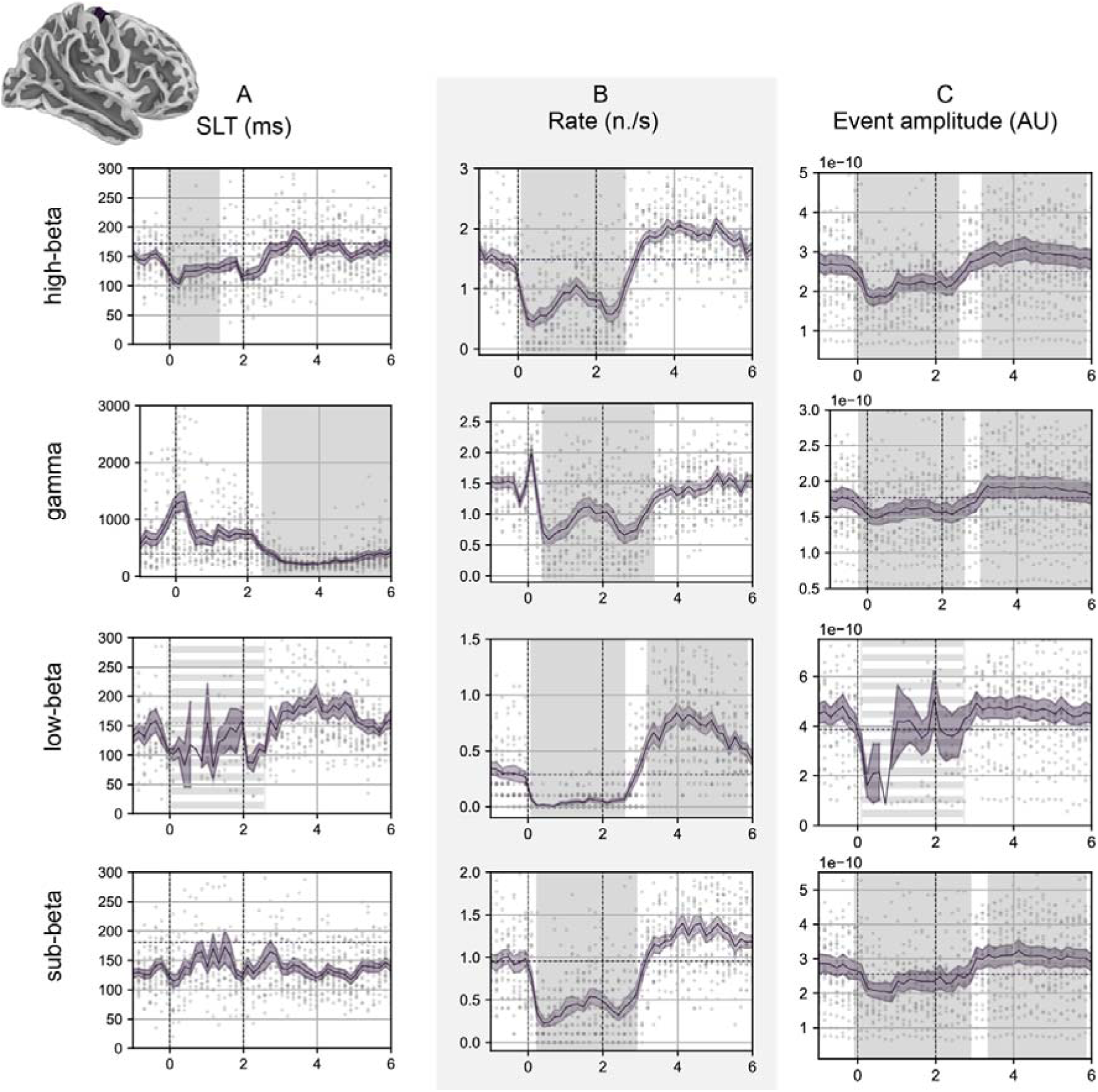
State characteristics’ modulation during motor task. **A.** State lifetimes, **B.** event rate and **C.** event amplitude modulation for the contralateral M1 parcel with peak FO modulation, visualized with sliding window average (window length 200 ms, overlap 50 ms). Individual dots indicate individual subjects’ values per time window, the black line and surrounding shading indicate the mean (line) +/- SEM (shading). Gray shaded areas indicate time windows differing significantly from baseline (non-parametric cluster-level pared t-test for non-spatial data implemented in MNE python, cluster threshold p < 0.05, significance threshold p< 0.001). The suppression interval of the low-beta state is marked with a vertical dashed line due to the scarcity of observed events, leading to decreasing reliability. Dashed horizontal red lines indicate resting state baseline level (resting eyes open), and vertical dashed black lines indicate movement initiation and termination cues (at 0 and 2 seconds). Note the differences in the ranges between different states (e.g., very long state lifetimes for gamma state).

**Supplementary figure 2:**
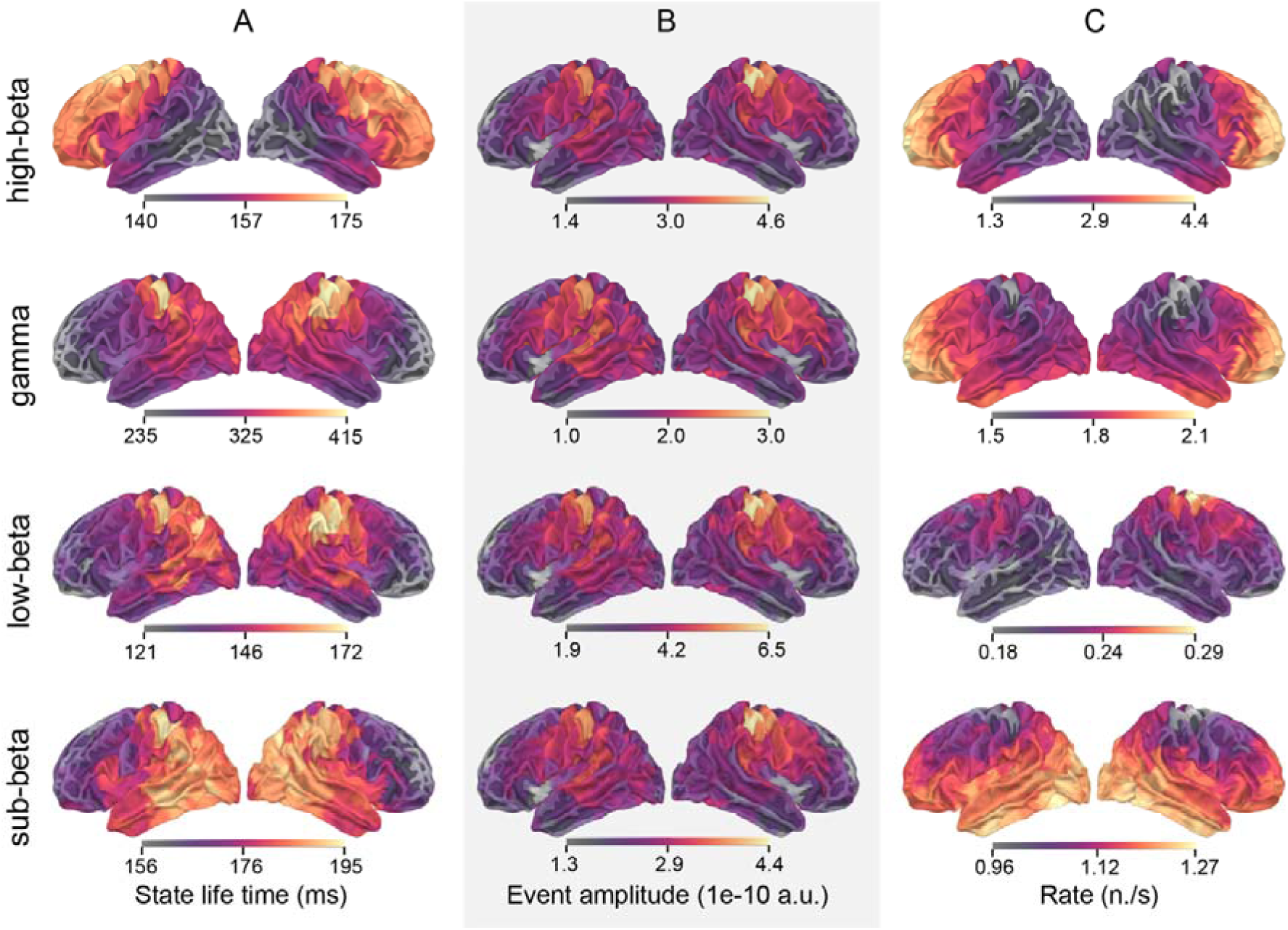
Topography of state characteristics across cortex. **A.** state lifetime, **B.** event amplitude and **C.** rate of occurrence. Note that while state lifetime and rate show state-specific spatial distribution maxima, amplitude maximum has the same location for all states. Also note that, e.g., for gamma state, the state lifetime spatial maximum differs from the rate maximum, i.e., the two carry partially orthogonal information. See Methods for details.

## Discussion

Using a Hidden Markov Model based approach, we can automatically decompose MEG activity in the 13-30 Hz (beta) range into a high- and a low-beta range state, enabling us to analyze the related beta event dynamics across the cortex at the parcel level. We show that both beta states occur throughout the cortex with distinct anatomical patterns and state characteristics: The low-beta state is a very rare, high amplitude event most likely to occur in sensorimotor areas, whereas the high-beta state is a common event with a maximum occurrence in the frontal areas. Both states show a suppression-rebound behavior during a motor task in a network consistent with previous work on motor task execution ^47^, with excellent signal-to-noise ratio. The pattern of suppression-rebound is different for the two beta states: While low-beta is suppressed entirely during movement execution, high-beta suppression is only partial, with periods of more pronounced suppression at initiation and termination of movement. Beta rebound is carried by the low-beta band and accompanied by markedly increased low-beta synchrony between brain areas. Movement termination is accompanied by a directed sequence of state transitions from the gamma/active through the high-beta to the low-beta oscillatory state, and then returns to the baseline state transition equilibrium. Information is also encoded via different state characteristics, such as event rate or lifetime, which can be modulated in orthogonal ways. Our findings shed light on the different roles of low- and high-beta band in motor processing and their dynamics, emphasizing the role of low-beta band in movement facilitation and termination.

Segregation of the beta band into low and high-beta band activity can often be seen already in individuals’ conventional PSD analysis ^35^ as separable peaks, and it has been described in the context of different brain processes with suggestions of preferential involvement of either band ^3,8,29^. Extending this, we here show a distinct functional profile for high versus low-beta band neuronal activity during movement. While low-beta is completely suppressed during sustained movement, high-beta modulation is dependent on task phase: A strong initial decrease with movement initiation is followed by a relative increase in high-beta state during static postural maintenance, as described previously for the beta band in general ^18–20,48^. High-beta is intermediate in contralateral motor areas during posture maintenance, whereas its levels return to baseline ipsilaterally, also suggesting that high-beta processing in contralateral motor cortex is involved in force maintenance. High-beta may thus correspond to, e.g., more distributed, cortical processing necessary for force maintenance and task monitoring during posture ^3,19^ and mediating cortico-muscular coherence ^20^, compatible also with the observed beta modulation during attentional processes and cue anticipation ^29,30^. The complete suppression of low-beta during movement, on the other hand, is compatible with a more general gating/releasing function for ‘putting thought into action’, compatible with previously postulated ‘anti-kinetic’ function of beta ^28,29^.

Beta rebound is dominated by an increase in low-beta event occurrence and associated with considerably increased synchronization. The pronounced synchrony across the network of low-beta events during post movement rebound could relate to reinforcing network connections after action execution, compatible with increased beta rebound during learning ^49^, reduction by error ^50^ and indexing confidence in outcomes ^25^. Using the proposed time- and frequency resolved analysis also allows investigation of transition dynamics between different oscillatory states during movement, showing that state transitions occur in a directed fashion, from active/gamma via high-beta to low-beta and back to baseline dynamics during the beta rebound. Movement initiation occurs via the high-beta state, while direct transitions between low-beta and gamma are absent, supporting low-beta’s previously proposed anti-kinetic role. In Parkinson’s disease (PD), low-beta band is frequently increased ^51^, and the lack of transitions between low-beta and active/gamma state observed in healthy subjects here suggest that a direct transition between both states does not normally occur, which may underline low-beta activity’s anti-kinetic effects. Levodopa increased the transition from low-beta state to other states in the subthalamic nucleus ^51^. Furthermore, voluntary movement initiation in PD can be improved, e.g., via visual cueing ^52^, which could be mediating the switch from low-beta to high-beta state, thus facilitating transition to active/gamma state.

The observed effects are unlikely to be contaminated by significant volume conduction between cortical areas. The task-related modulation of the states shows clear, spatially non-contiguous modulated areas in ipsilateral M1/S1 (iM1/S1), as well as in the prefrontal cortex; the modulation observed in iM1 differs qualitatively from contralateral M1 (cM1). Furthermore, the modulation in cM1 is clearly localized in the hand area, and the network of modulated anatomical areas coincides with typical sensorimotor network areas. High resolution MEG studies have also shown beta waves to “travel” along the cortex ^53^.

Overall, the proposed approach offers a novel framework for studying whole-brain beta band activity in relation to behavior in a time- and frequency resolved manner, applicable not only to sensorimotor processing but also potentially for studying beta dynamics in, e.g., cognitive processes. Our results add to the understanding of brain oscillatory dynamics during movement initiation, maintenance, and termination, and reconcile previously incongruent findings about the role of beta in movement-related processing. This can have implications, e.g., for adaptive, closed loop deep brain stimulation approaches targeting beta band activity in PD or brain computer interfaces. Furthermore, the increased information obtained from whole-brain electrophysiological data obtained via simultaneously time- and frequency-resolved analysis can benefit electrophysiology-based biomarker development.

## Materials and methods

### Subjects and MEG recordings

MEG measurements were performed in a magnetically shielded room (Imedco AG, Hägendorf, Switzerland) with a 306-channel Vectorview neuromagnetometer (Elekta Neuromag TRIUX, MEGIN Ltd, Helsinki, Finland) consisting of 204 planar gradiometers and 102 magnetometers, at the BioMag Laboratory at Helsinki University Hospital. Five head position indicator (HPI) coils were attached to the face and scalp, and their locations were digitised with a 3-D digitiser pen (Fastrak, Polhemus, US). The 3-D digitiser was also used to digitise the subject’s head shape and three anatomical landmarks (the nasion and the right and left preauricular points). Spontaneous cortical activity was recorded with a 1 kHz sampling rate, continuous head position monitoring (cHPI) and band-pass filtering at 0.03-330 Hz.

MEG data from a total of 42 healthy subjects (27 males and 15 females, age mean +/- STD 45.7 +/- 14.8 years, range 18-78 years, 37 right/3 left-handed and two ambidextrous) were used in the study. All subjects participated in a resting eyes open (restEO) measurement, and n=22 subjects (12 males and 10 females, age mean +/- STD 48.5 +/- 15.2 years, range 18-78 years, 20 right-handed and 2 ambidextrous) additionally participated in a left-hand motor task (leftmotor). During the resting state measurement, subjects were resting with their eyes open. In the motor task, subjects were instructed to squeeze a ball with their left hand after an auditory cue und maintain the squeeze until a second auditory cue at two seconds was presented, followed by 5 seconds rest (total trial duration 7s).

### MEG data pre-processing

For suppressing external artefacts, MEG data were pre-processed using the temporally extended signal space separation method (tSSS) ^54^, implemented in the MaxFilter software (MEGIN Oy, Helsinki, Finland, version 2.2.15). Head movements were compensated based on the simultaneous cHPI recordings. Individual head positions were kept to avoid additional transformation-related noise in the source reconstruction.

Further signal processing was done using MNE-python version 1.7 ^42,43^. Heart-related and eye movement artefacts remaining after tSSS were removed using the FastICA algorithm ^55^ based on visual inspection of the resulting component time series and topography. Additionally, segments with muscle artefact were visually annotated as bad data segments and removed. ICA and annotations utilized data that were filtered to include frequencies ranging from 1 Hz to 40 Hz.

The data was then filtered to beta-band between 13 and 30 Hz with a one-pass, zero-phase, non-causal band-pass FIR (finite impulse response) filter. The lower and upper transition bandwidths were 3.25 Hz and 7.50 Hz with cutoff frequencies at 11.38 Hz and 33.75 Hz. The filter length was 1017 samples, using a Hamming window with 0.0194 passband ripple and 53 dB stopband attenuation. Finally, the MEG data was downsampled to 200 Hz.

### MEG data reconstruction to parcel-level source space

The bandpass-filtered pre-processed MEG data was reconstructed from the sensor to the source space using FreeSurfer’s fsaverage MRI atlas ^56^. The fsaverage MRIs were individually scaled based on the digitised head shape points, and the MEG-MRI transformation was then done with the scaled fsaverage MRI. The boundary element model (BEM) needed for the forward model was created using the watershed algorithm ^57^ in MNE-Python. The source space, describing the positions of the candidate source locations, was set up with 5120 source points. The orientation of the sources in the forward solution was fixed to be perpendicular to the cortex ^58^. Inverse modelling was done using MNEs, with signal-to-noise ratio value of 1.0, loose orientation constraint = 0, and depth weighting of = 0.8. Since the data was a continuous time series, the noise covariance was calculated using empty room recordings acquired on the measurement day after applying tSSS and filtering (13-30 Hz).

The dimensionality of the source reconstructed MEG signals (5120 source points) was reduced to 448 cortical parcel time series using a subdivided aparc parcellation (aparc_sub, available in MNE) ^44^ and principal component analysis of the timeseries as implemented in MNE-Python (mode=’pca_flip’; ^42^). Here, principal component analysis finds the dominant temporal pattern, and signal cancellation is prevented by flipping signals whose orientation differed from the dominant direction within the parcel ^59^. The resulting parcel time series were used for further analysis.

### Hidden Markov Models for time-resolved analysis of oscillatory states in MEG data

The Hidden Markov Model (HMM) approach segments the MEG time series data into a sequence of a finite number of states ^40,46^. In practice, the HMM assigns each data point a probability of belonging into each state, as each data point is assumed to be drawn from a probabilistic observation model. The family of distributions (a Gaussian distribution here) for the observation models is the same for all states. We used the Time-Delay Embedded (TDE) observation model ^46^, which instead of only focusing on one observation *Y_t_* at the point *t*, looks at the points from *t*-*L* to *t*+ *L* (here 85 ms window). This allows estimating the relevant frequency as part of the of the state parameters by utilizing an autocovariance matrix, which characterizes the spectral properties of the signal within the 85-millisecond time window. Therefore, the observations (here MEG data points) become *Y_t_* = (*y_t_*_-*L*_, …, *y*, …, *y_t_*_+*L*_) and the TDE observation model becomes

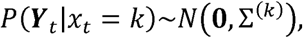

where each state *k* has individual observation model parameters, mean and autocovariance, which are the mean of the time series and the linear relations across regions and time points within a time window around a given time point *t* ^60^.

Since we wanted to characterize the oscillatory features independently of the signal’s amplitude properties, we pinned the mean to zero. After an initial testing phase, we set the number of states to be *k*= 4 and the lag parameter to be *L*= 8. Assignment to a certain state is done following the order-one Markovian, meaning that the probabilities of states being active at time *t* are only affected by the state that was active at time *t*-1. The estimation of the posterior distribution parameters is inferred using variational Bayes and stochastic inference ^40,61^. Given MEG data as input, each time point is assigned probabilities of belonging to each of the states, which are called the state time courses. Based on the state time courses, binary time courses for each state can be derived (so-called Viterbi path). Additionally, the power spectral densities (PSD) of the states are determined for each state separately using the multitaper method ^40^.

We first fit the HMM uniformly for all subjects, i.e., on a group-level: *N* subjects’ *p* parcel-level time courses with *t* timepoints are concatenated, resulting in data with the dimensionality ℝ^1×*Npt*^ which is used to train a group-level model. Compared to the more typical input format of ℝ*^p^*^×*Nt*^, which assumes that the state frequency is the same across all brain areas, this allows later estimating individual versions of the parameters of interest both across subjects and across parcels. The group-level model’s states are common for each subject, meaning that the states’ probability distribution parameters are obtained uniformly for all subjects, i.e., assuming that similar activity falls into the same state in all subjects. This is necessary to ensure that the states are comparable between subjects and parcels. We then re-fit the HMM separately to each individual and parcel (input dimensions ℝ^1×*t*^) to obtain individual- and parcel-specific versions of the states (so called ‘dual’ estimation) ^41^.

This preserves the general definition of the group-level states but adjusts the group states to better fit each individual and brain area. To avoid overfitting, the group level model was only trained on resting (restEO) data from n=36 subjects and then used to dual-estimate all subjects’ and parcels’ individual HMM for both the resting as well as the motor task data. We used the Matlab version of the HMM MAR toolbox for data analysis ^40^, but there is now an alternative Python implementation ^64^.

### Oscillatory state characteristics

For further analysis, we used the probability time courses and binary state time courses (i.e., Viterbi paths) from the individually estimated HMMs to obtain fractional occupancy (*FO*), as well as additional descriptive event characteristics, namely state lifetimes (*SLT*), event amplitudes (*EA*) and events per second (*rate*) for each state (**figure 1, left bottom panel**). The events’ state lifetimes, i.e., duration of individual state visits to each state, were calculated taking the mean of all state visits longer than 50 ms for each state, assuming that shorter lifetimes reflect spurious switches. Fractional occupancy, i.e., the share of time spent in one state out of the total time, was calculated by dividing the number of time points spent in a state by the total number of time points in the time series. Rate was derived by dividing the total number of state visits per state by the total duration of the signal for the resting data, giving an estimate of events per second. Event amplitude was calculated as follows: To obtain the amplitude envelope, the original 13-30 Hz band-filtered parcel MEG time series was decomposed with a set of complex Morlet wavelets within the frequency range with 1 Hz resolution and *n_cycles = frequency/*2. Then, the mean of the absolute value of the signal’s time-frequency representation was taken over the beta range (13-30 Hz) to obtain its amplitude envelope. Finally, the mean of all timepoints within a state visit exceeding the 75^th^ percentile of values was calculated for each individual state visit.

### Oscillatory state dynamics

Next, we analysed state characteristic modulation during the motor task. We first calculated task-evoked FOs for each state over time by averaging the state probability (FO) time courses across trial epochs, resulting in a task-related FO modulation. Epochs were averaged over -1 to 6 s relative to the start cue, -1 to 0 seconds were used as baseline FO. Spatio-anatomical patterns of task-related modulation were also analysed (**figure 3C**, see also *Statistical testing*). To calculate the other task-related state characteristics modulation (state lifetime, event amplitude and rate), we used a sliding window average (window length 200 ms, 50 ms overlap).

### Oscillatory state synchronization

State synchronization between different states was analysed using the binary state time courses (Viterbi path) and was carried out for low and high-beta states. For both resting (restEO) and task-related (leftmotor) data, event coincidence was assessed by counting, for pairs of 2 parcels, the time points at which the state of interest occurs simultaneously in both parcels, and dividing that by the duration of the observed data segment, i.e., the fraction of coincident time points with respect to the total number of time points (similarly to Tinkhauser et al. 2018 ^62^). The event coincidences between pairs of two parcels were then divided by the respective measure from resting state, to enable comparison between high and low-beta states and reduce the impact of FO differences between the states. This was done for the suppression (0-3 s) and rebound (3-6 s) periods.

This coincidence measure was calculated for all possible aparc-sub parcel combinations within the top 30% of parcels of the spatiotemporal clusters with significant FO modulation during the task (‘evoked FO’ clusters, see *Statistical testing*). For ease of interpretation, we visualized the original high-dimensional coincidences on a coarser parcellation, the Desikan-Killiany atlas ^63^, using the parcels subsuming the most significant areas of the evoked FO clusters (visualized in **figure 3B**, ‘precentral-lh’, ‘postcentral-lh’, ‘superiorparietal-rh’, ‘superiorfrontal-rh’, and ‘rostralmiddlefrontal-rh’ contralaterally, and ‘precentral-rh’ and ‘postcentral-rh’ ipsilaterally). The transformation from the sub-parcellation (aparc_sub) to the coarser Desikan-Killiany parcellation was done by taking the maximum connection among all aparc_sub parcel pairs within each Desikan-Killiany pair.

### State transition dynamics

Finally, we investigated state transition dynamics during the motor task in the contralateral M1 parcel showing most significant FO modulation in the spatiotemporal clustering test (**figure 3C**). All state transitions throughout the time series can be grouped according to state pair and transitions direction (n=12 possible pairs). The directional transitions between the state pair (e.g., transitions from low-beta to high-beta) were visualized during the motor task using sliding window average (window length 100 ms, 25 ms overlap) and compared to baseline to detect significant transition changes (see *Statistical testing* and **figure 4)**.

### Statistical testing

To investigate spatio-temporal patterns in the task-related activity, a two-sided non-parametric cluster-level paired t-test for spatio-temporal data, as implemented in MNE-Python, was performed on the source space data. This identified the statistically significant spatio-temporal FO clusters for different states (number of permutations 1024, degrees of freedom *n_subjects* - 1, cluster forming threshold *p* < 0.001, significance threshold *p* < 0.001, see **figure 3C**). Here, the multiple comparison problem is addressed with a cluster-level permutation across space and time.

The same test for non-spatial data was used to test if the FO evoked curves differed from baseline (**see figure 2E and 2F**). An exploratory analysis was additionally done for 7 pre-defined areas showing significant spatio-temporal FO modulation clusters (visualized in **figure 3B**, ‘precentral-lh’, ‘postcentral-lh’, ‘superiorparietal-rh’, ‘superiorfrontal-rh’, and ‘rostralmiddlefrontal-rh’ contralaterally, and ‘precentral-rh’ and ‘postcentral-rh’ ipsilaterally, **figure 3A).** For contralateral M1 (one parcel in ‘precentral-lh’ area), we also investigated whether low-beta FO was significantly different from zero, as well as whether there were differences between low-beta and high-beta state, by scaling the curve with the mean of the baseline (-1 to 0 s) and with baseline set to zero. The cluster threshold was set to *p* < 0.001, and the significance threshold *p* < 0.001 for all analyses. The number of permutations was 1024 and degrees of freedom *n_subjects* - 1.

A two-sided, non-parametric cluster-level paired t-test for non-spatial data was used to test significance of other time-dependent state characteristics (amplitude, state lifetime, event rate). While the FO-evoked curve has a value for each time point and subject, features based on single events are much sparser and different from high-frequency sampled ‘conventional’ MEG timeseries data, affecting statistical power.

Thus, more lenient thresholds were chosen for this analysis. The rate, state lifetime and event amplitude, were tested with respect to the baseline with a cluster threshold of *p* < 0.05, a significance threshold of *p* < 0.001, number of permutations 1024 and degrees of freedom *n_subjects* - 1 (**figure 5**). State transition dynamics were tested using MNE’s two-sided non-parametric cluster-level paired t-test with the same thresholds and parameters (**figure 4**). Since we observed very few transitions between gamma state and low-beta state and vice versa, we additionally tested whether the observed transition occurrences match the transitions expected based purely on each state’s fractional occupancy, using a chi-square distribution test (chi-square goodness of fit test, significance threshold used was *p* < 0.001). Suppression (0-3 s) and rebound (3-6 s) periods were tested separately. Observed transitions were obtained by taking the sum of all observed transitions over a period per state pair, and classifying them (for low-beta – gamma state transitions, there are transitions between the two states in either directions, and transitions to other states, as follows: low-beta -> gamma, low-beta -> other; gamma -> low-beta, gamma -> other). Expected transitions were calculated by dividing the observed total transitions according to the proportions expected from the fractional occupancy mean across subjects during the analysis period (suppression, rebound).

To test for significant differences in state synchronization between high and low-beta states, we used the synchronization values of all included parcel pairs (here *aparc_sub* parcellation) and treated those as one group (not corrected for multiple comparison). We compared rest vs. task during suppression (separately for low and high-beta states), rest vs. task during rebound (separately for low and high-beta states), and high-beta vs. low-beta (task values relative to rest, tested separately for suppression and rebound). Here we used a two-sided Mann-Whitney U-test with significance threshold of *p* < 0.001.

## Funding and acknowledgements

CA was funded by a Carlsberg Foundation Visiting Postdoctoral Fellowship at the University of Oxford (CF23-1716). DV was supported by a Novo Nordisk Foundation Emerging Investigator Fellowship (NNF19OC-0054895) and an ERC Starting Grant (ERC-StG-2019–850404). HR received funding from the Research Council of Finland (grant numbers 355409 and 321460). KAMP was in part supported by the Research Council of Finland (grant number 350242), the Sigrid Juselius Foundation, the Finnish Medical Foundation, and a government research grant (valtion tutkimusraha).

## Author roles

SK: conceptualization, formal analysis, data curation, methodology, software, visualization, writing-original draft preparation

CA: methodology, writing – review & editing

LN: data curation, investigation, writing – review & editing

DV: methodology, writing – review & editing

HR: investigation, resources, writing – review & editing

AP: conceptualization, methodology, investigation, funding acquisition, project administration, supervision, visualization, writing-original draft preparation

